# Prediction of Bovine coronavirus immune epitopes using selection and phylogenetic analysis

**DOI:** 10.1101/2023.09.14.557617

**Authors:** Tristan Russell, Jose Maria Lozano, Cailyn Challinor, Gerald Barry

**Affiliations:** UCD School of Veterinary Medicine, University College Dublin, Ireland; Central Veterinary Research Laboratory, Department of Agriculture, Food and the Marine, Celbridge, Ireland

**Keywords:** Bovine coronavirus, Immune epitopes, Selective Pressure, Phylogenetics, Ireland

## Abstract

*Bovine betacoronavirus* (BoCoV) is a pneumoenteric pathogen of cattle, which is closely related to Human coronavirus OC43. Vaccines are administered to protect against diseases caused by BoCoV, but knowledge gaps exist with regard to correlates of protection and the effect of immune evasion on driving evolution. In this study, immune epitopes were predicted for the BoCoV structural proteins including spike and haemagglutinin esterase (HE) and these predictions were supported through targeted gene sequencing of Irish clinical isolates and selective pressure analysis. Increased prevalence of diversifying selection and amino acid changes in some predicted immune epitopes suggests immune escape is selecting for non-synonymous mutations arising in these regions. Selection analysis and sequencing provided increased support for nAb epitopes compared to other predicted immune epitopes suggesting nAbs are an important arm of the immune response to BoCoV. Phylogenetic analysis of spike and HE sequences showed Irish isolates from this study were in the European clade except for one HE sequence that sat in the Asian/American clade, while the spike gene of this sample was in the European clade. Recombination between a European and an Asian/American isolate could give rise to such a sequence and recombination breakpoints that were detected at the 3’ end of HE and 5’ end of spike would produce such a sequence. This study presents evidence showing pressure to evade the nAb response is contributing to BoCoV evolution and for the first time, sequenced an isolate likely derived from a recombination event between European and Asian/American strains.

## Introduction

*Bovine betacoronavirus* (BoCoV) is a pneumoenteric pathogen of *Bos taurus* though closely related viruses have been isolated from several other ruminant species and some non-ruminant mammals [1]. In cattle it causes winter dysentery, calf diarrhoea [2] and is associated with bovine respiratory disease [3]. In Ireland a retrospective study published in 2014 found that of the five viruses associated with the respiratory disease complex included in the screen, BoCoV was the most frequently detected in calves with respiratory disease in Ireland [4]. BoCoV is also frequently detected in pneumonia cases as part of co-infections [5].

As a member of the *Embecovirus* subgenus, BoCoV encodes five structural proteins including four conserved across the *Coronaviridae* family – spike, nucleocapsid, membrane and small envelope – as well as a surface protein unique to this *Embecovirus* subgenus – haemagglutinin esterase (HE). HE is a receptor degrading enzyme, which facilitates virus entry and release by targeting the 9-O-acetylated sialic acid residues required for BoCoV entry [6]. It has previously been shown infection-induced humoral immune responses against these proteins can protect cattle from disease [1, 7, 8], while some antibodies generated in response to vaccine can recognise antigens derived from clinical isolates [9]. However, it has been shown calf BoCoV antibody titres do not correlate with protection from respiratory disease when dams received a vaccine designed to protect against enteric BoCoV disease [10, 11].

Our current understanding of BoCoV immune epitopes, correlates of protection, and the effect of immune evasion on BoCoV evolution is limited [1]. To further understand which viral proteins are targeted by the immune system and to assess the contribution of immune evasion pressure on protein sequence variation, predicted immune epitopes of BoCoV structural proteins were combined with selective pressure analysis in this study. There are already full-genome sequences of 24 BoCoV isolated from cattle in 2019/20 from across Ireland available publicly [12] as well as ten S1 domain and one full-length spike sequences from isolates collected in the province of Munster in 2010/11 [13]. Here the spike and HE genes of 29 Irish clinical isolates were sequenced and compared with other BoCoV isolates obtained nationally and internationally. Phylogenetic analysis of the spike and HE sequences identified residues where amino acid changes have occurred compared to a commonly used vaccine strain. We present evidence of selective pressure on residues associated with predicted immune epitopes suggesting that vaccination is driving variation in these genes, and this may lead to reduced efficacy of currently used vaccines.

## Materials and Methods

### Samples and Detection of Bovine coronavirus

Nasopharyngeal samples collected from 29 cattle displaying symptoms of respiratory illness during 2022/23 in Ireland were submitted to the Department of Agriculture, Food and the Marine’s Virus reference laboratory (Table 1). Samples were collected as part of normal veterinary diagnostic practices. These samples were then tested for the presence of respiratory pathogens by qPCR. Those positive for BoCoV were identified using a previously developed TaqMan qPCR protocol [14]. Sequencing of positive samples was carried out as approved by the University College Dublin animal research ethics committee, approval code AREC-E-22-18-Barry.

**Table 1:** Sample details and GenBank accession numbers. Abbreviations: ND, no disease information; NA, no animal information; BHV-1/4, *Bovine alphaherpesvirus-1/4*; BRSV, *Bovine respiratory syncytial virus*; PIV-3, *Parainfluenza virus-3*; d, days old; w, weeks old; m, months old; y, years old.

| Sample no. | GenBank Accession | Location (county) | Date | Pathology | Host |
| --- | --- | --- | --- | --- | --- |
| 1 | OR271239 | Galway | 30/03/2022 | ND | NA |
| 2 | OR271240 | Waterford | 11/05/2022 | ND | NA |
| 3 | OR271241 | Sligo | 27/04/2022 | Dead, pneumonia, *Histophilus somni* and *Mannheimia haemolytica* coinfection | 3 w |
| 4 | OR271242 | Laois | 08/04/2022 | ND | 2 m |
| 5 | OR271243 | Laois | 08/04/2022 | ND | 2 m |
| 6 | OR271244 | Wexford | 22/03/2022 | Enteric symptoms, *Escherichia coli* co-infection | 5 w |
| 7 | OR271245 | Wexford | 21/03/2022 | Enteric and respiratory symptoms, *E. coli* co-infection | 10 d |
| 8 | OR271246 | Kilkenny | 16/03/2022 | ND | Calves (pooled) |
| 9 | OR271247 | Kilkenny | 14/03/2022 | Pneumonia and enteritis, BHV-1 co-infection | 4 w |
| 10 | OR271248 | Cavan | 23/02/2022 | *Pasteurella multocida* and *M. haemolytica* co-infection | 4 animals (pooled) |
| 11 | OR271249 | Donegal | 15/02/2022 | Pneumonia | 10 m |
| 12 | OR271250 | Wexford | 11/03/2022 | Pneumonia, other bacterial respiratory pathogens isolated | 2 w |
| 13 | OR271251 | Wexford | 11/03/2022 | Pneumonia, other bacterial respiratory pathogens isolated | 2 w |
| 14 | OR271252 | Sligo | 06/01/2022 | ND | Weanling |
| 15 | OR271253 | Laois | 08/04/2022 | ND | < 1 m |
| 17 | OR271254 | Wicklow | 23/05/2022 | Pneumonia, BHV-4, *H. somni* and *M. haemolytica* co-infections | 3 m |
| 116 | OR271255 | Westmeath | 2022 | Dead | 1 y male |
| 117 | OR271256 | Offaly | 2022 | Pneumonia | 8 m |
| 119 | OR271257 | Meath | 2022 | Pneumonia | 6 w |
| 123 | PP156983 | Carlow | 04/11/2022 | Pneumonia, BRSV co-infection | 9 m |
| 124 | PP156990 | Kilkenny | 03/11/2022 | Respiratory symptoms | 2 y female |
| 125 | PP156992 | Kilkenny | 06/12/2022 | Respiratory and enteric symptoms, PIV-3 co-infection | 8 m |
| 126 | PP156984 | Cork | 14/12/2022 | Dead, respiratory and enteric symptoms, BRSV co-infection | 6 m |
| 127 | PP156985 | Galway | 21/11/2022 | Dead, pneumonia, BRSV co-infection | 7 m |
| 128 | PP156986 | Longford | 02/12/2022 | Respiratory symptoms, PIV-3/BRSV co-infections | 5 w |
| 129 | PP156987 | Carlow | 26/10/2022 | Respiratory and enteric symptoms, PIV-3 co-infection | 5 m |
| 131 | PP156988 | Wexford | 10/11/2022 | Pneumonia, enteric symptoms, BRSV co-infection | 10 m |
| 132 | PP156991 | Tipperary | 06/01/2023 | Enteric symptoms | 10 m female |
| 133 | PP156989 | Carlow | 07/11/2022 | Dead, pneumonia | 8 m |

### Recombination and Selection analysis

BoCoV spike, HE, nucleocapsid, membrane and small envelope coding sequences were downloaded from the GenBank database (https://www.ncbi.nlm.nih.gov/, Table S1). Identical and incomplete sequences were removed from datasets. Codon alignments for each gene were obtained using the Clustal Omega alignment algorithm [15, 16] in Jalview version 2 [17–19]. Stop codons were deleted manually. Recombination breakpoints of each gene were predicted using Genetic Algorithm for Recombination Detection (GARD) [20] through the data monkey webserver (datamonkey.org) [21]. Sequences arising through recombination were removed from alignments, then mixed effects model of evolution (MEME) [22] and fast unconstrained bayesian approximation (FUBAR) [23] methods were used through the data monkey web server [21] to infer site-specific selective pressures. MEME predicted sites under diversifying episodic selection and FUBAR estimated synonymous and non- synonymous mutation rates to infer pervasive selective pressure.

### Prediction of immune epitopes

Immune epitopes for BoCoV structural proteins were predicted based on three previous studies. Firstly, BoCoV spike and HE Major Histocompatibility Complex-I (MHC-I, also known as bovine leukocyte antigen-I) and B cell receptor (BCR) immune epitopes were predicted bioinformatically [24]. The second two methods mapped immune epitopes identified for *Human betacoronavirus OC43* (HCoV OC43) protein onto BoCoV orthologues because, depending on the variants selected for comparison, spike, HE and NCP shared approximately 91%, 94% and 97% identity at amino acid level. Also, the two Embecovirus species are closely related, with evidence suggesting HCoV OC43 evolved from BoCoV following a relatively recent zoonotic event [25]. MHC-II (also known as bovine leukocyte antigen-II) immune epitopes identified for HCoV OC43 by co-immunoprecipitation (co-IP) of MHC-II molecules [26] were mapped onto BoCoV spike, HE, nucleocapsid and small envelope . HCoV OC43 spike residues shown to interact with neutralising antibodies (nAb) through structural analysis or shown to be escape mutants following passage in the presence of antibodies [27] were also mapped onto BoCoV spike.

### Protein Structures

BoCoV spike structure was predicted using SWISS-MODEL (https://swissmodel.expasy.org/) with the BoCoV Mebus spike amino acid sequence as input [28]. BoCoV HE structure had been determined by x-ray crystallography [29] and deposited on the protein data bank (https://www.rcsb.org/). Spike and HE structures were visualised and annotated in PyMOL [30].

### Sequencing spike and haemagglutinin esterase

RNA from samples positive for BoCoV was converted to cDNA using the SuperScript III Reverse Transcriptase kit (Invitrogen) following the manufacturer’s protocol. cDNA was used as a template for PCR (Phusion HF kit (New England Biolabs)) with primers designed to amplify the spike and HE genes (Table 2). PCR products from reactions producing a specific fragment underwent PCR clean-up (Qiagen) and, if multiple fragments were detected, gel purification (Qiagen) from a 0.8% agarose gel was carried out. Purified fragments were sent for Sanger sequencing using appropriate forward and reverse primers (Eurofins Genomics).

**Table 2:**
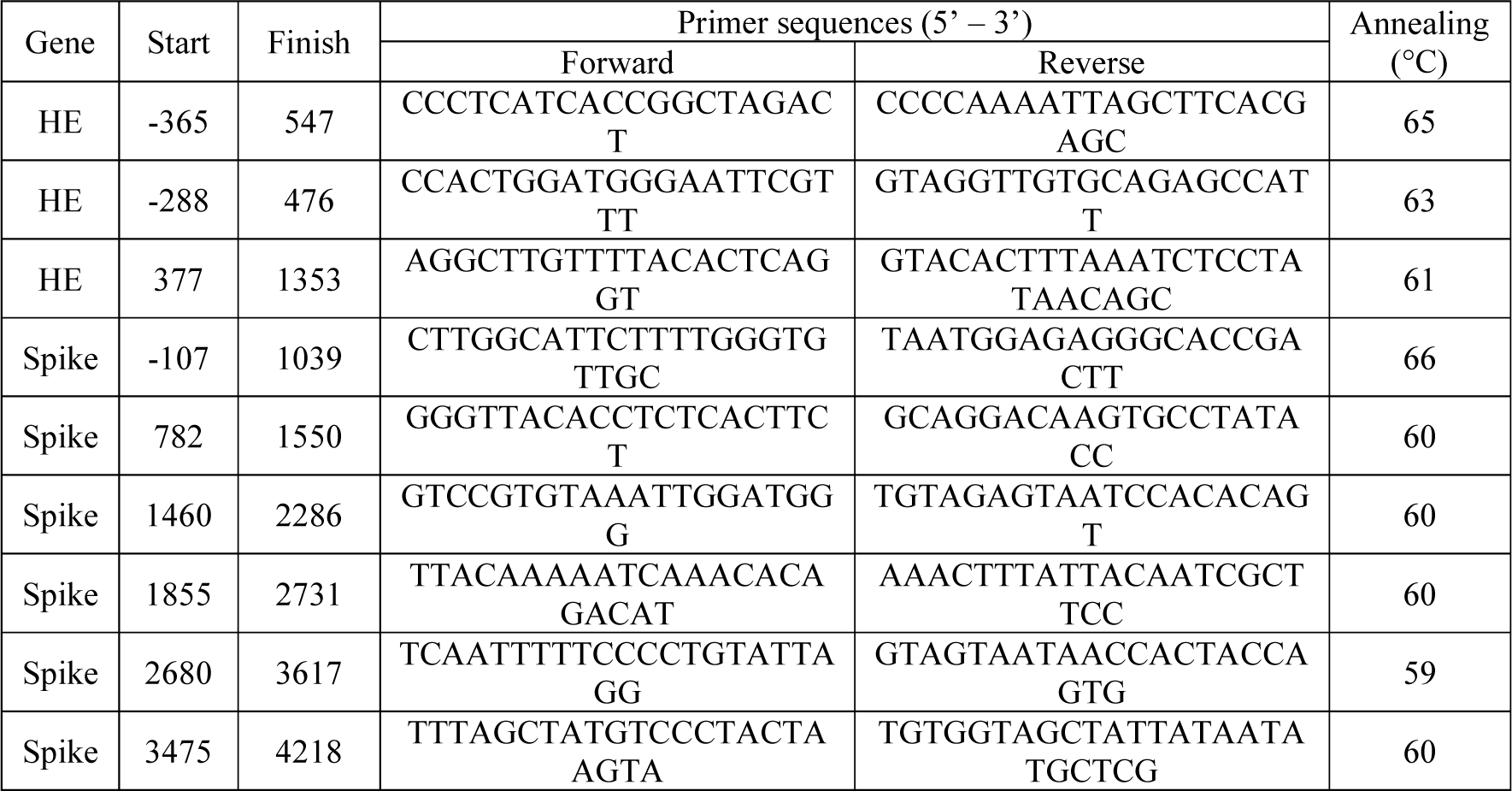
Primer pairs used to amplify regions of HE and spike genes. Start and finish positions of the amplicons are provided relative to the first nucleotide of the HE or spike genes. Primers were designed for this study or taken from. [31, 32].

### Phylogenetic analysis

Reads obtained from Sanger sequencing were trimmed to remove low quality base calls. Reads from the same sample were aligned to BoCoV spike and HE references using the Clustal Omega alignment algorithm [15, 16] in Jalview [17–19]. References were selected based on the closest NCBI Blast hits for individual sequences from the same sample. Sequences were deposited on GenBank (Table 1).

Nucleic acid sequences were converted to amino acid sequences then phylogenetic trees were generated using the and neighbour-joining method [33] in Seaview version 5 [34]. Trees were annotated and images generated in the Interactive Tree of Life web server [35].

## Results

### Selection Analysis of BoCoV structural proteins

Regions of proteins under diversifying selection could represent immune epitopes, several MHC-I, MHC-II, BCR and nAb epitopes within BoCoV structural proteins were predicted as described in the Materials and Methods and are presented here in Table 3. Site-specific selective pressures were then inferred using FUBAR and MEME algorithms (Figure 1). Prior to running these, GARD analysis was used to identify and remove sequences derived from recombination. Recombinant sequences were detected in the spike gene alignment, but not genes of other BoCoV structural proteins. Spike (Figure 1A) and HE (Figure 1B) were under increased diversifying selection compared to nucleocapsid (Figure 1C), membrane (Figure 1D) and small envelope (Figure 1E) proteins as demonstrated by the increased proportion of codons predicted to be under diversifying selection for spike and HE compared to other structural proteins (Table 4). There was increased abundance of codons under purifying selection within functional domains such as the fusion peptides of spike (Figure 1A); the esterase domains of HE (Figure 1B); the RNA binding and dimerisation domains of nucleocapsid (Figure 1C); and the transmembrane domains of membrane (Figure 1D) and small envelope (Figure 1E). Receptor binding domains (RBD) are involved in protein-protein interactions and often contain nAb epitopes so, selection at spike and HE RBD was of particular interest. Increased selection was observed for the RBD of spike with 9.46% codons within RBD under diversifying selection compared to 5.35% for the whole gene and 47.97% codons within the RBD under purifying selection compared to 41.76% for the whole gene (Figure 2A). There was increased diversifying selection for the HE RBD with 4.90% codons within the RBD under diversifying selection compared to 4.44% for the whole gene but decreased purifying selection as 19.58% of codons within the RBD were under purifying selection compared to 22.66% for the whole gene (Figure 2B).

**Figure 1:**
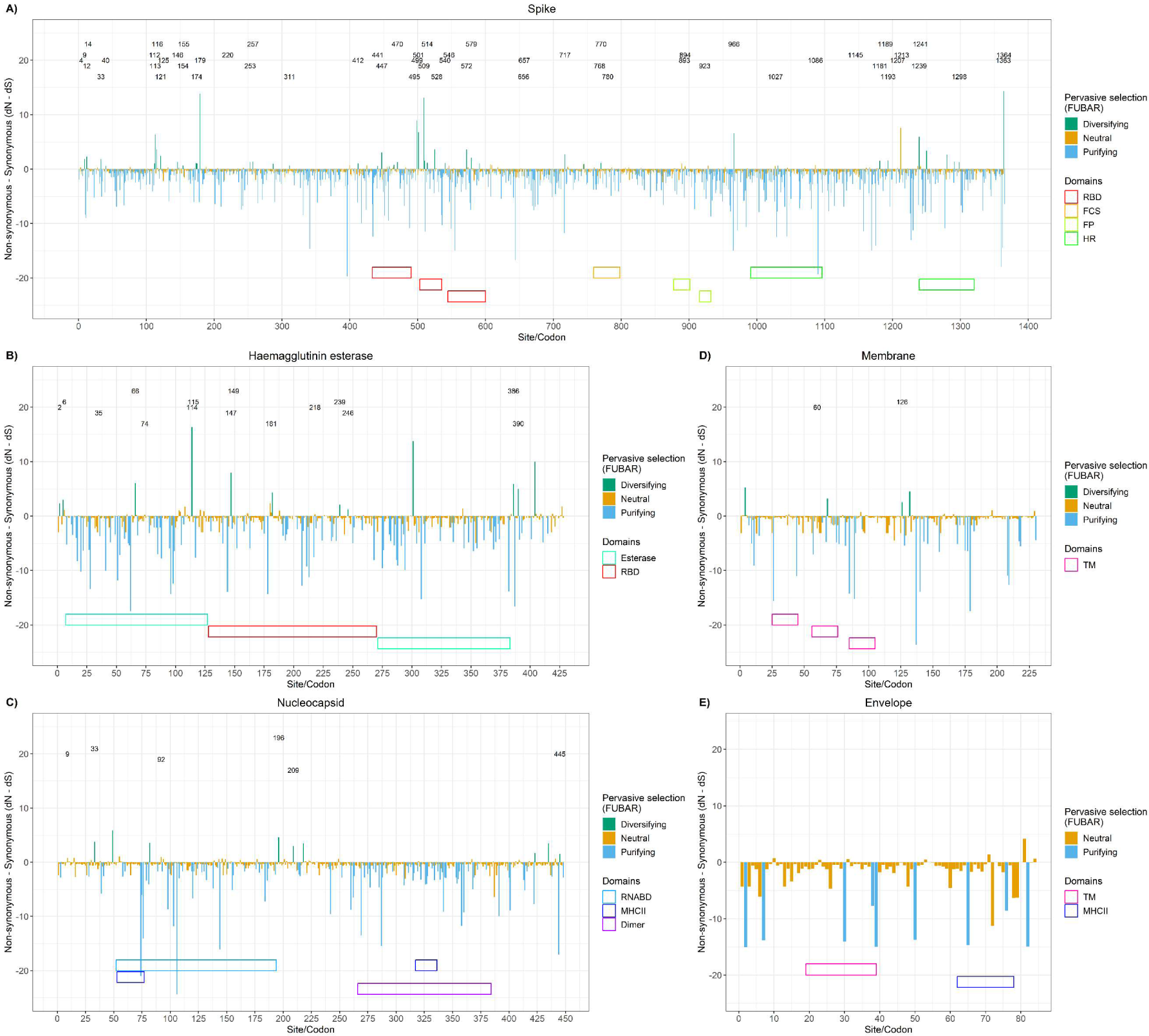
Selection analysis of structural BoCoV proteins. Bar plots show the results of selection analysis with FUBAR and annotated codons were predicted to be under diversifying selection using MEME for spike (A), HE (B), nucleocapsid (C), membrane (D) and small envelope (E) proteins. Locations of structural and functional domains were annotated at the bottom of plots. Abbreviations: Dimer, interaction interface for dimerisation; FCS, Furin Cleavage Site; FP, fusion peptide; HR, heptad repeat; RBD, receptor binding domain; RNABD, RNA binding domain; and TM, transmembrane domain.

**Figure 2:**
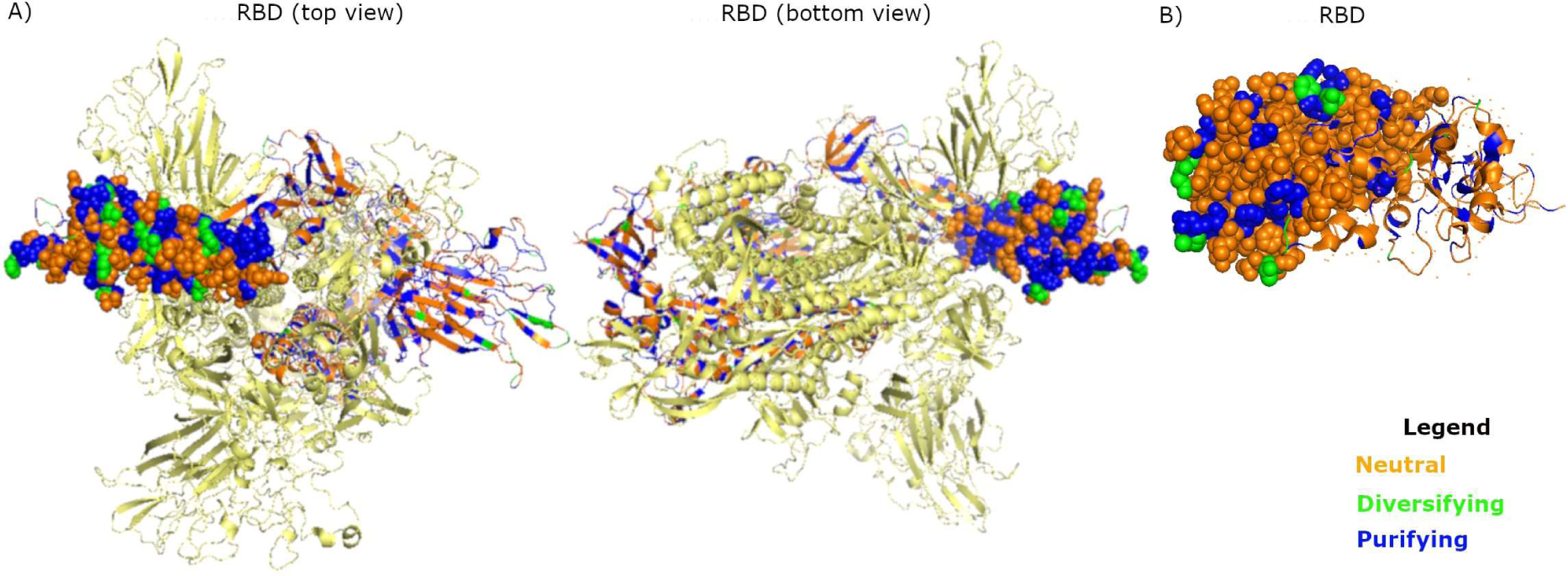
Residues in the RBD of spike (A) and HE (B) shown as spheres with residues coloured based on selection analysis of corresponding codons.

**Table 3:**
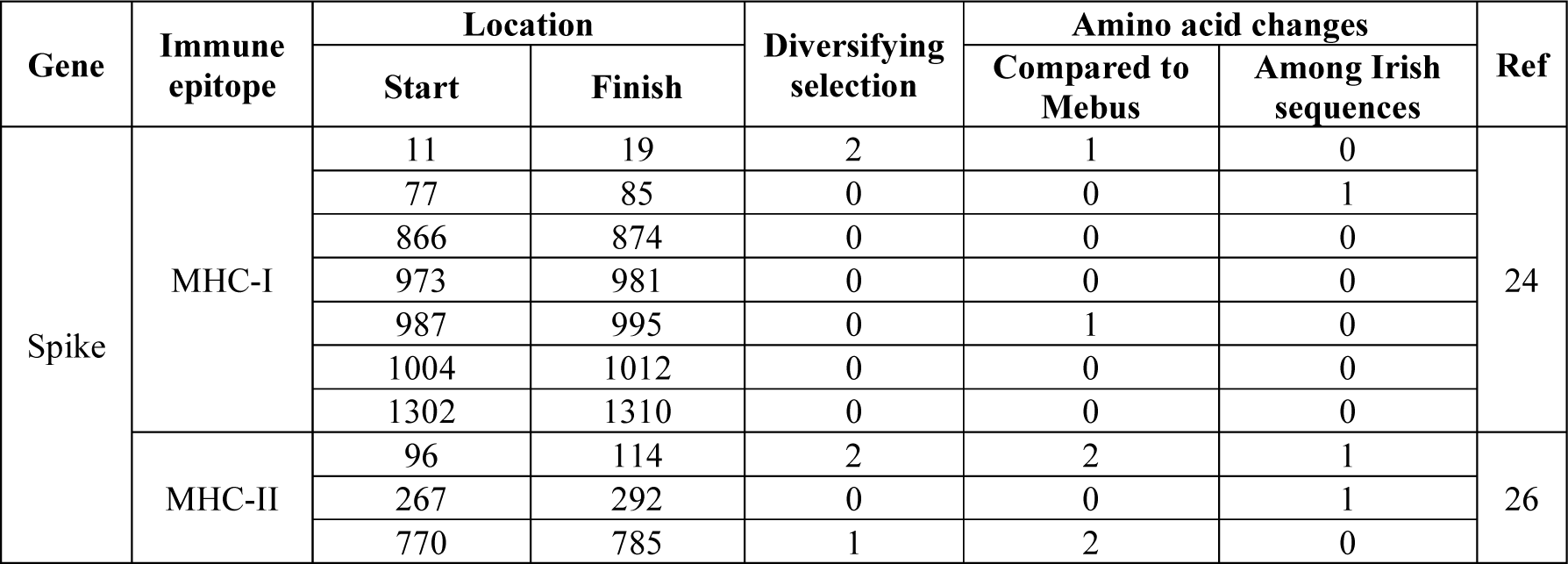
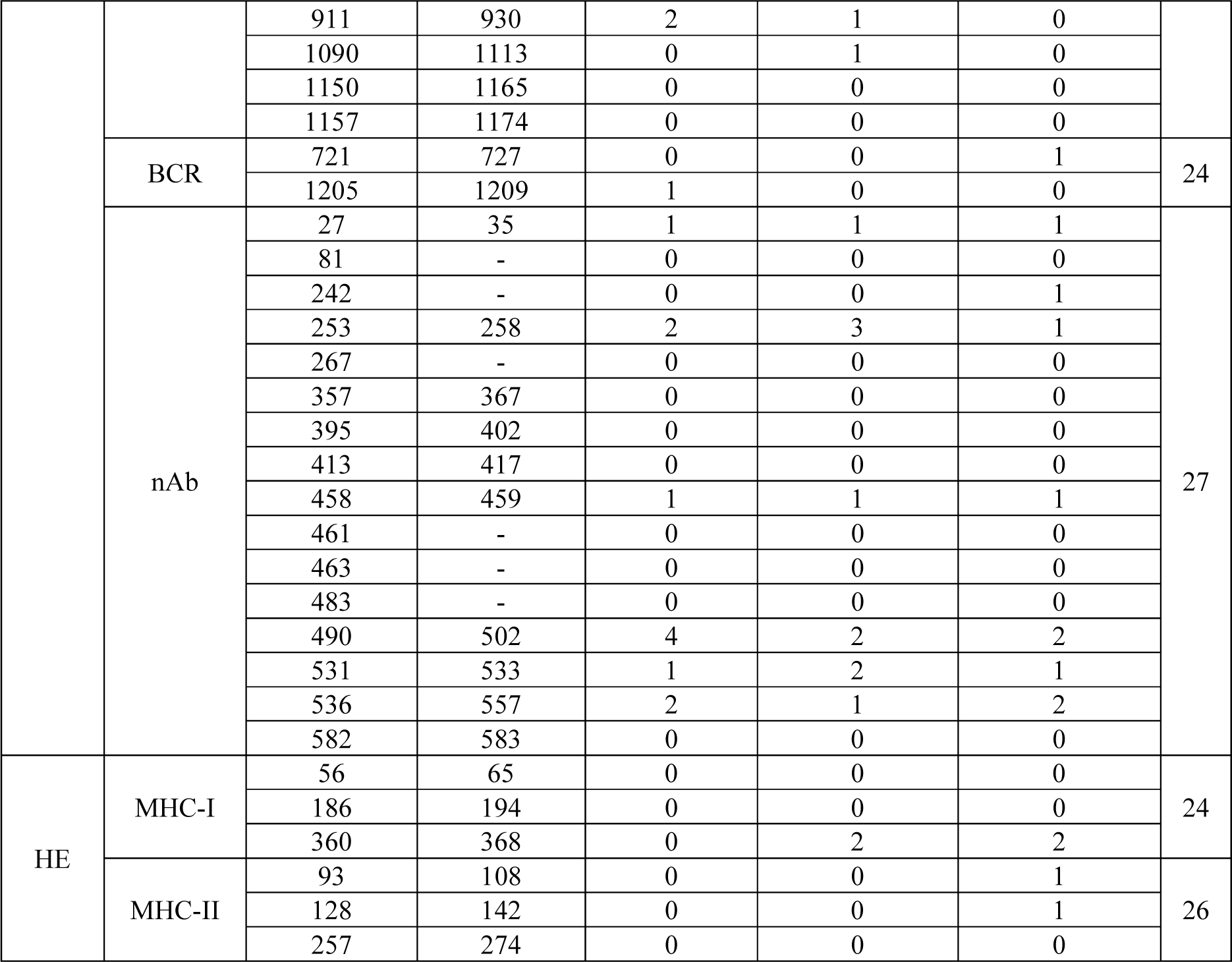
Predicted immune epitopes and overlap with codons under diversifying selection and residues where amino acid changes were observed.

**Table 4:**
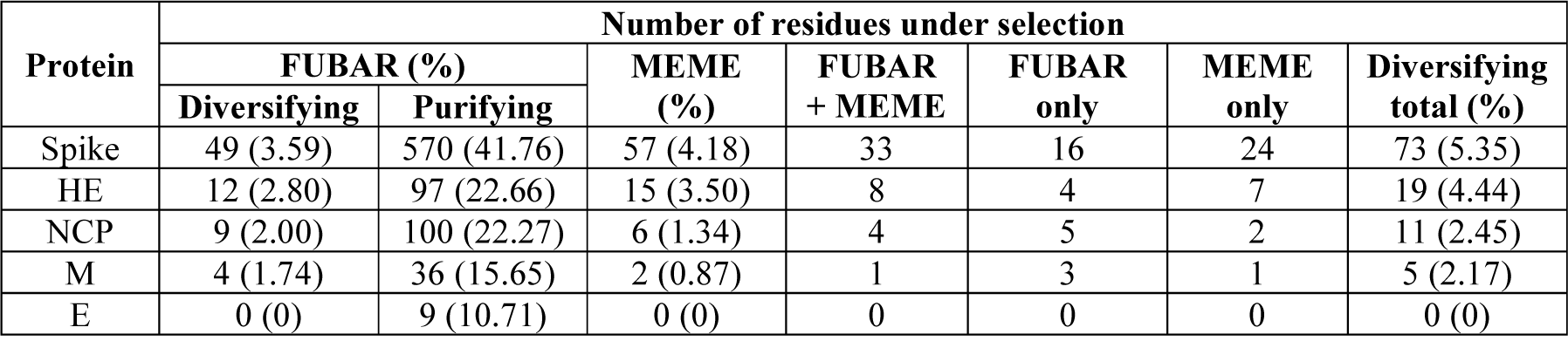
Selection analysis of structural proteins. Number and proportion of codons found to be under selective pressure for each BoCoV structural protein as determined by FUBAR and MEME.

| Protein | Number of residues under selection |  |  |  |  |  |  |
| --- | --- | --- | --- | --- | --- | --- | --- |
|  | FUBAR (%) |  | MEME (%) | FUBAR + MEME | FUBAR only | MEME only | Diversifying total (%) |
|  | Diversifying | Purifying |  |  |  |  |  |
| Spike | 49 (3.59) | 570 (41.76) | 57 (4.18) | 33 | 16 | 24 | 73 (5.35) |
| HE | 12 (2.80) | 97 (22.66) | 15 (3.50) | 8 | 4 | 7 | 19 (4.44) |
| NCP | 9 (2.00) | 100 (22.27) | 6 (1.34) | 4 | 5 | 2 | 11 (2.45) |
| M | 4 (1.74) | 36 (15.65) | 2 (0.87) | 1 | 3 | 1 | 5 (2.17) |
| E | 0 (0) | 9 (10.71) | 0 (0) | 0 | 0 | 0 | 0 (0) |

Immune escape provides a selective advantage for amino acid changes occurring in immune epitopes so selection within predicted BCR, MHC-I, MHC-II and nAb immune epitopes was assessed (Table 3). Spike codons under diversifying selection overlapped one of seven spike MHC-I epitopes and one of two spike BCR epitope predicted computationally (Figure 3A). Three of seven spike MHC-II immune epitopes predicted by co-IP overlapped codons under diversifying selection (Figure 3A). Of the 87 residues predicted to contribute to nAb recognition 12.64% were under diversifying selection compared to 5.35% codons of the whole spike gene (Figure 3A). Diversifying selection was not observed for any predicted HE MHC-I or MHC-II immune epitopes (Figure 3B), nor the two predicted nucleocapsid and one predicted small envelope MHC-II immune epitopes.

**Figure 3:**
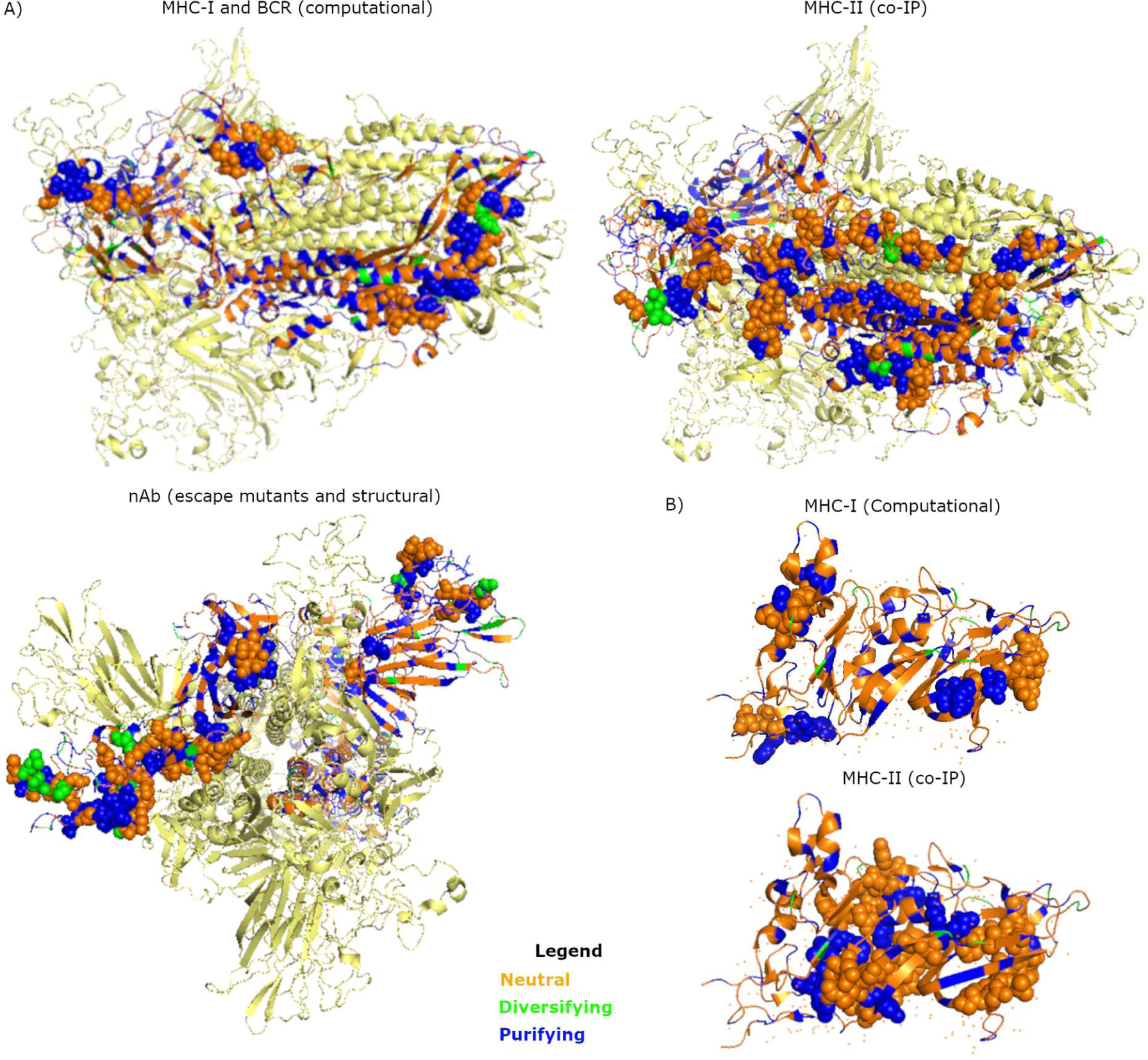
Spike trimer and HE structures with residues coloured based on results of FUBAR and MEME. Only one spike monomer was coloured. A) Spike immune epitopes predicted computationally, by co-IP of HCoV OC43 and by structural analysis of HCoV OC43 nAb-protein interactions were shown as spheres. B) Predicted HE MHC-II and MHC-I immune epitopes shown as spheres. Abbreviations: BCR, B cell receptor; co-IP, co-immunoprecipitation; MHC-I, major histocompatibility complex I; MHC-II, major histocompatibility complex II; and nAb, neutralising antibody.

### Sequencing and phylogenetic analysis of BoCoV spike and HE

To compare currently circulating Irish isolates with older Irish isolates, international isolates and vaccine strains, the spike and HE genes of 29 BoCoV clinical isolates from cattle presenting with respiratory and sometimes enteric symptoms in Ireland were sequenced. Full-length spike and HE sequences were obtained for 27 and 22 isolates, respectively. All spike sequences, except Sample 129, fell into one of two clades (Figure 4A), while most HE sequences were split between two clades with Sample 117 and Sample 133 the only exceptions (Figure 4B). Most samples sat in the same clade of the spike and HE trees but Sample 123 and Sample 128 switched clade, while Sample 117 and Sample 133 were both in one of the main clades in the spike tree (Figure 4A) but formed outgroups in the HE tree (Figure 4B). Differences in the spike and HE phylogenetic trees suggests recombination between BoCoV circulating in Ireland. GARD analysis of a HE-spike nucleotide alignment of sequences obtained in this study, starting from the start codon of HE to the stop codon of spike, detected seven recombination breakpoints at positions 360, 1260, 2023, 2562, 3475, 4612 and 4915 (Supplementary Figure S1). The first two overlap the HE gene and the next five overlap the spike gene in positions 733, 1272, 2185, 3322 and 3625. As previously shown, spike sequences of natural isolates collected globally predominantly diverged based on location with one clade of Asian and American isolates and a second clade of European isolates containing the Irish sequences from this study (Figure 4C). Some spike sequences from this study were in a subclade containing other Irish isolates collected in 2019/20, while most were in a subclade of French isolates from 2013/14 (Figure 4C). Sample 129 branched off on its own and was closest to a subclade of Swedish isolates from 2010. An enteric Irish isolate from 2011 diverged from other sequenced BoCoV isolates in its own subclade (Figure 4C). A tree of global HE sequences also mainly diverged based on location with one clade of Asian and American isolates and a clade of European isolates (Figure 4D). Some HE sequences from this study shared a subclade with Irish isolates collected in 2019/20, most formed a separate subclade with Albanian isolates collected in 2023, while Sample 117 was in the Asian/American clade and closest to enteric Chinese isolates from 2018 and 2020 (Figure 4D). GARD analysis of the spike-HE alignment of all GenBank sequences and sequences from this study detected three recombination breakpoints at positions 1549, 3073 and 4567, which all overlapped the spike gene at positions 247, 1771 and 3265 (Supplementary Figure S2).

**Figure 4:**
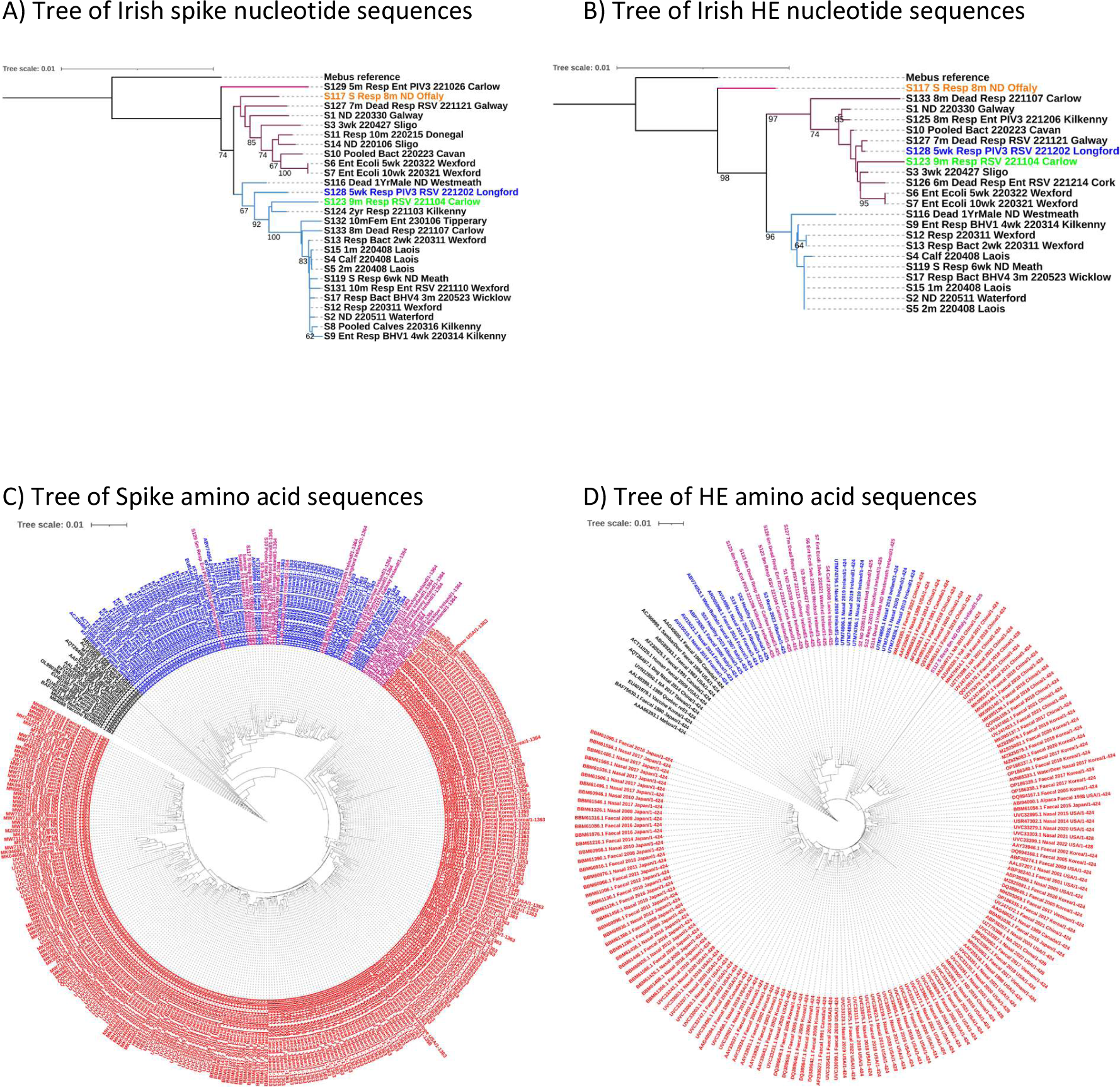
Phylogenetic trees of BoCoV spike and HE generated using the neighbour joining method and rooted using the Mebus strain as an outgroup. A) Nucleotide sequences of spike for isolates from this study and the Mebus strain with different coloured branches indicating two main clades. B) Same as A), but for HE. The colours of branches for each clade match in A) and B), while isolates with coloured labels switched clades or formed their own outgroup. C) All unique Spike amino acid sequences downloaded from GenBank, with label colours indicating location of isolation with Asian/American in red, European in blue and old isolates/vaccine strains in black. Sequences from this study are shown in pink. D) The same as C), but for HE.

Consensus sequences of spike and HE obtained in this study were translated and compared with BoCoV Mebus because this strain was used to design an inactivated vaccine often used in Ireland. Pressure to evade vaccine-mediated immunity would provide a selective pressure for amino acid changes occurring in immune epitopes of natural isolates so they diverge from vaccine strains. Compared to Mebus, there were amino acid changes in 48 (3.52%) spike residues and 9 (2.12%) HE residues of the Irish consensus sequences. As prior infection with natural isolates can generate an effective immune response against future infection, there could also be selection for non-synonymous mutations among Irish BoCoV isolates. Among the spike sequences obtained in this study there were amino acid changes in 61 (4.47%) residues, while in HE there were amino acid changes in 23 (5.42%) residues. Residues containing amino acid changes were grouped based on those different from Mebus (Mebus mutations) and those where variation was observed when comparing sequences from this study (Ireland mutations) (Figure 5). Non-synonymous Mebus mutations overlapped two MHC-I epitopes (3.17% of residues predicted to be MHC-I epitopes), four MHC-II epitopes (4.32%) and 10/87 residues predicted to be involved in nAb recognition (11.49%) in spike (Figure 5A). Ireland mutations overlapped with one BCR (8.33%), one MHC-I epitope (1.59%), two MHC-II epitopes (1.44%) and 9/87 residues predicted to be involved in nAb recognition in spike (10.34%) (Figure 5A). One predicted HE MHC-I epitope overlapped with Mebus and Ireland mutations (7.14%) while two predicted MHC-II epitopes overlapped with Ireland mutations only (6.12%) (Figure 5B).

**Figure 5:**
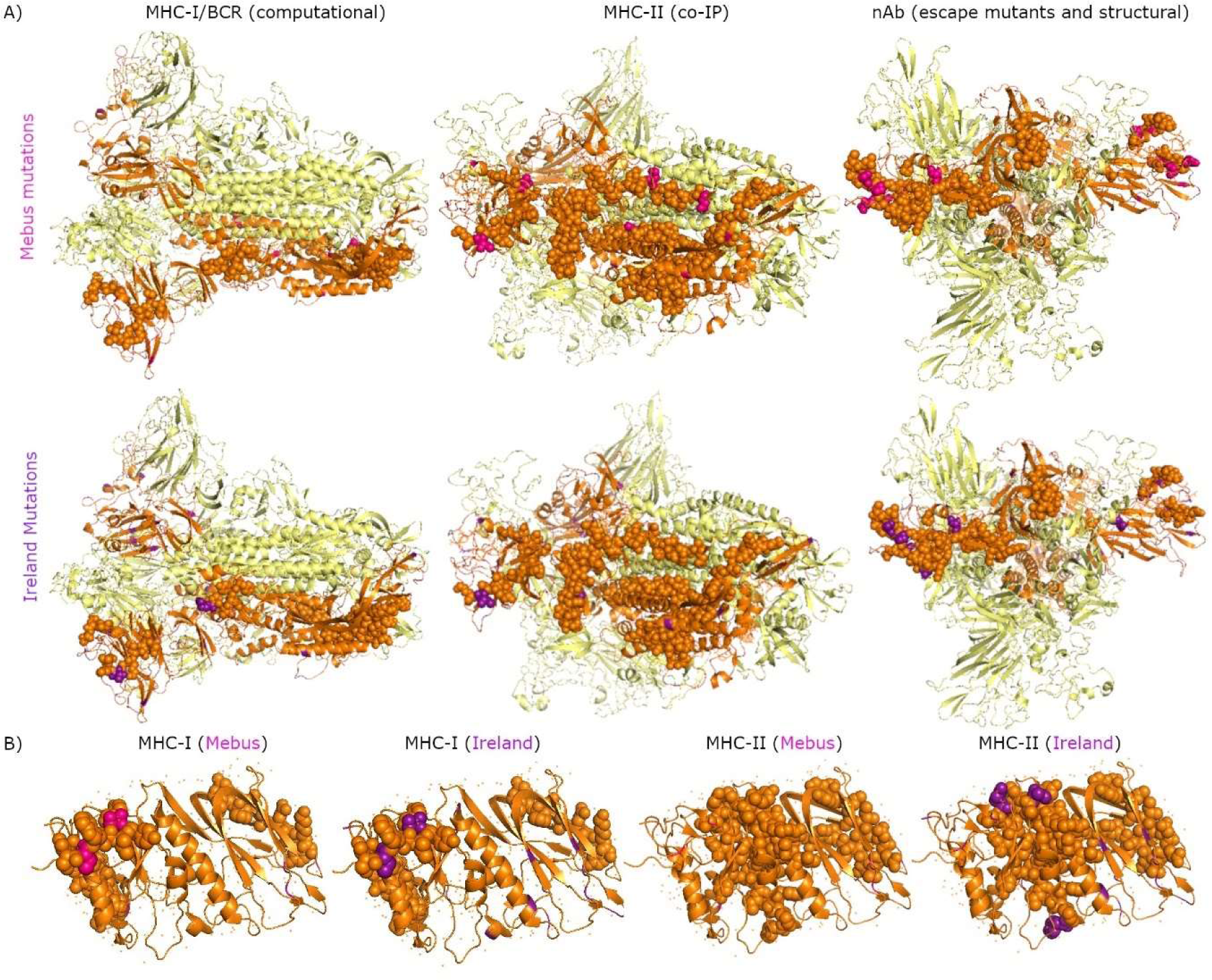
Overlap between residues with amino acid changes and predicted immune epitopes. A) The spike trimer is presented as a cartoon, with predicted immune epitopes presented as spheres on one spike monomer in orange. Mebus mutations are shown in pink and Ireland mutations in purple. B) Same as A) but for HE.

## Discussion

Site specific selective pressures can be used to predict residues important to protein structure and function because these residues are more likely to be under purifying selection. As expected, there was overlap between residues under purifying selection and several annotated structural and functional domains of BoCoV proteins. The high level of selection observed for spike RBS is expected because its interaction with the entry receptor would drive purifying selection [36, 37], while RBD often contain nAb epitopes, which would drive diversifying selection [38, 39]. Several residues within the HE RBD were also under selection with an increased prevalence of diversifying selection compared to the rest of the protein. The high level of purifying selection in the esterase domains, which have catalytic activity and do not overlap the RBD, means there was not an increased prevalence of purifying selection for HE RBD compared to the rest of the protein.

Three studies were used to predict immune epitopes of structural BoCoV proteins. BoCoV spike and HE MHC-I and BCR immune epitopes were predicted bioinformatically [24]. HCoV OC43 spike, HE, NCP and envelope MHC-II epitopes were identified by co-IP [26], then mapped onto BoCoV orthologues. Residues and domains involved in nAb recognition of HCoV spike were determined by structural analysis and identification of escape mutants [27], then mapped onto BoCoV spike. Pressure to evade the immune response is one driver of diversifying selection in *Coronaviridae* [36, 40], although there are other drivers of diversifying selection, such as adaptation to a new host [41]. Therefore, selective pressures acting on residues along BoCoV structural proteins were analysed to provide additional evidence as to the true nature of the predicted immune epitopes with the assumption that some diversifying selection would be driven by immune escape. The proportion of codons under diversifying selection along the whole genes of spike and HE were increased compared to other structural proteins suggesting they are more likely to contain immune epitopes. There was no overlap between residues under diversifying selection and predicted HE immune epitopes but 5.96% of residues in predicted spike immune epitopes were under diversifying selection compared to 5.35% of residues for the full-length protein. There were more residues under diversifying selection within nAb immune epitopes than other predicted immune epitopes. There was also overlap between predicted immune epitopes and amino acid changes within the sequences from this study suggesting immune escape could be selecting for these changes. Once again, an increased proportion of amino acid changes were observed in nAb epitopes compared to other predicted immune epitopes implying there is a selective advantage for non-synonymous mutations in regions recognised by nAbs. Selection for amino acid changes in predicted nAb epitopes suggests nAbs are involved in an effective BoCoV immune response to BoCoV. In the case of MHC-I, reduced support for these epitopes by could be due to downregulation of the MHC-I presentation by BoCoV. HCoV OC43 [26] and other *Coronaviridae* species have been shown to suppress MHC-I antigen presentation [42, 43]. Therefore, there would be no selective advantage to non-synonymous mutations in these regions.

The COVID-19 pandemic has shown that administration of a vaccine can drive diversification and reduce the protection offered by the vaccine to natural isolates. Widespread administration of a vaccine means most the population harbours an effective immune response against a particular version of the virus, which selects for amino acid changes occurring in immune epitopes. Updating the vaccine to resemble the clinical isolates more closely is one strategy to maintain effectiveness of the vaccine. Natural isolates evolving away from the Mebus vaccine strain appears to be occurring in Ireland because amino acid changes were observed in 48 (3.52%) and 9 (2.12%) spike and HE residues, respectively. These amino acid changes may also occur in regions important to immune recognition because 13 changes in spike and 4 changes in HE overlap with predicted immune epitopes. It is currently unclear whether this evolution of the circulating BoCoV strains has reduced the effectiveness of the vaccine, but the rate of continual infection and disease associated with BoCoV suggests it may be contributing. Designing a subunit vaccine based on currently circulating Irish clinical isolates of BoCoV could generate a more protective vaccine for cattle based in Ireland.

Almost all spike sequences from this study were in one of two clades suggesting most isolates descended from two distinct introductions of the virus into Ireland or diverged along two separate branches following a single introduction. Sample 129 formed an outgroup on its own and was the only isolate not grouped into the two main clades. This divergence from other sequences could be driven by adaptation to a unique environment or be the result of a third introduction of BoCoV into Ireland, which is less frequently detected. Less frequent detection could be caused by reduced sampling frequency associated with reduced pathogenicity, or it could be caused by reduced prevalence, which could indicate a recent introduction into Ireland. Another possibility is that this variant has a selective disadvantage causing it to be outcompeted by other variants thus reducing its prevalence. Relatedness of Sample 129 with other isolates from this study was only based on the spike gene because full-length HE sequence was not obtainable. The HE of most sequences from this study fell into one of two clades, which predominantly overlapped with the two main clades of the spike tree. Two isolates, namely Sample 123 and Sample 128, switched clade suggesting recombination had occurred around the 3’ end of the HE gene or 5’ end of the spike gene, which was supported by GARD analysis showing breakpoints around both regions. Sample 117 also no longer clustered with either of the two main clades of the phylogenetic tree. Interestingly, when phylogenetically classified Sample 117 sits in the Asian/American clade, which could indicate recombination between a European isolate and an Asian/American isolate. This would require co-infection of an animal with both strains. Import and export of cattle into and from Ireland is rare [44], especially with other continents, so one possibility is that the recombination event occurred elsewhere in Europe and the animal was then brought into Ireland. Alternatively, an Asian/American strain of BoCoV could have been brought to Ireland by another mammalian species and the recombination event occurred in Ireland.

## Conclusion

Phylogenetic analysis of sequences from this study suggested there have been at least two introductions of BoCoV into Ireland, and for the first time, sequence obtained from a European isolate has been shown to sit in the Asian/American clade along with evidence of recombination between European and Asian/American isolates. This study has predicted several BoCoV immune epitopes and corroborated several of those in spike and HE with selection analysis and amino acid changes observed from sequence data suggesting some of the predicted immune epitopes could be recognised by the bovine immune response. Overlap between predicted immune epitopes and amino acid changes observed when comparing the Mebus vaccine strain with clinical isolates could reduce the protection offered by Mebus- based vaccines. It is recommended that studies to assess the impact of this virus evolution on the effectiveness of currently used vaccines is assessed to identify whether vaccines require updating.

## Acknowledgement

We thank the team of people in the Department of Agriculture Food and the Marine diagnostic laboratory for screening bovine samples for the presence of BoCoV by PCR.

## Supplementary Information

**Figure S1:**
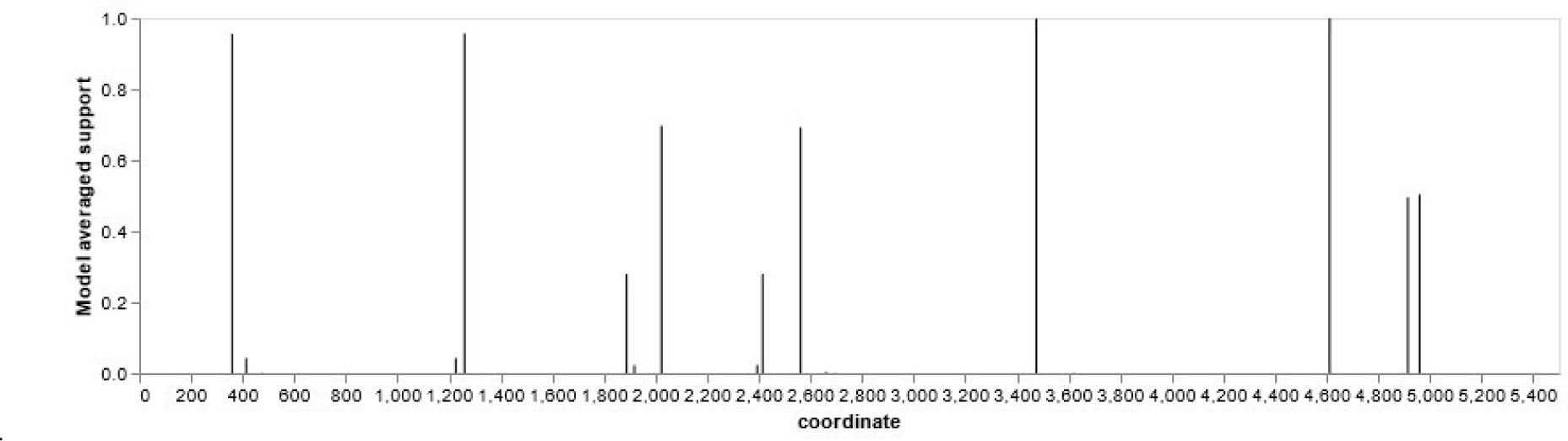
Recombination breakpoints detected in HE-S nucleotide alignment of sequences obtained in this study. The 20 full-length spike-HE sequences were aligned using Clustal Omega then recombination breakpoints detected using GARD.

**Figure S2:**
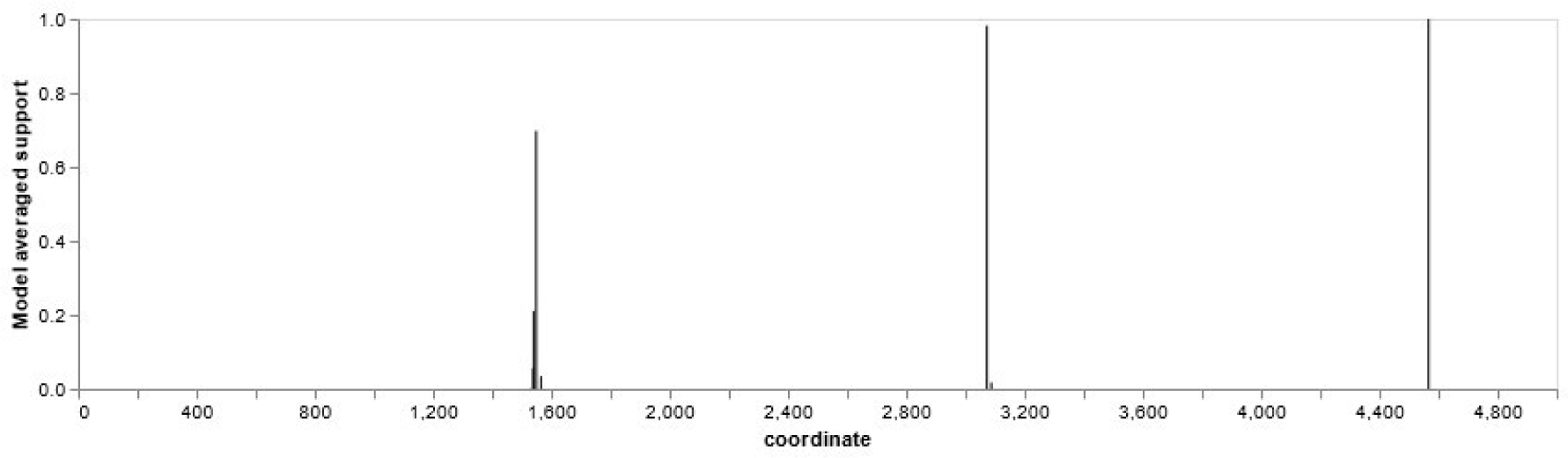
Recombination breakpoints detected in HE-S nucleotide alignment of GenBank sequences and sequences obtained in this study. Sequences were aligned using Clustal Omega then recombination breakpoints detected using GARD.

**Table S1:**
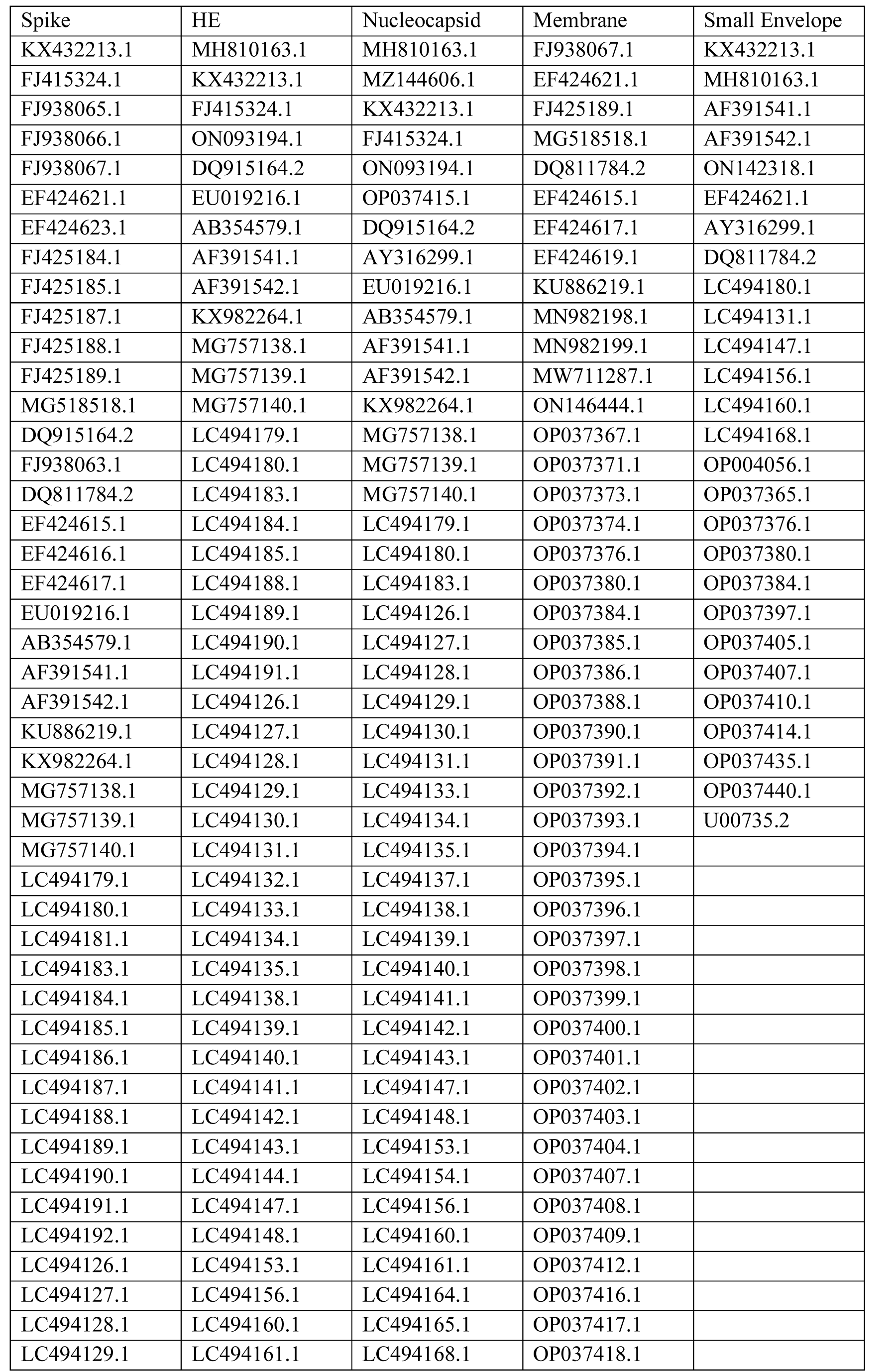
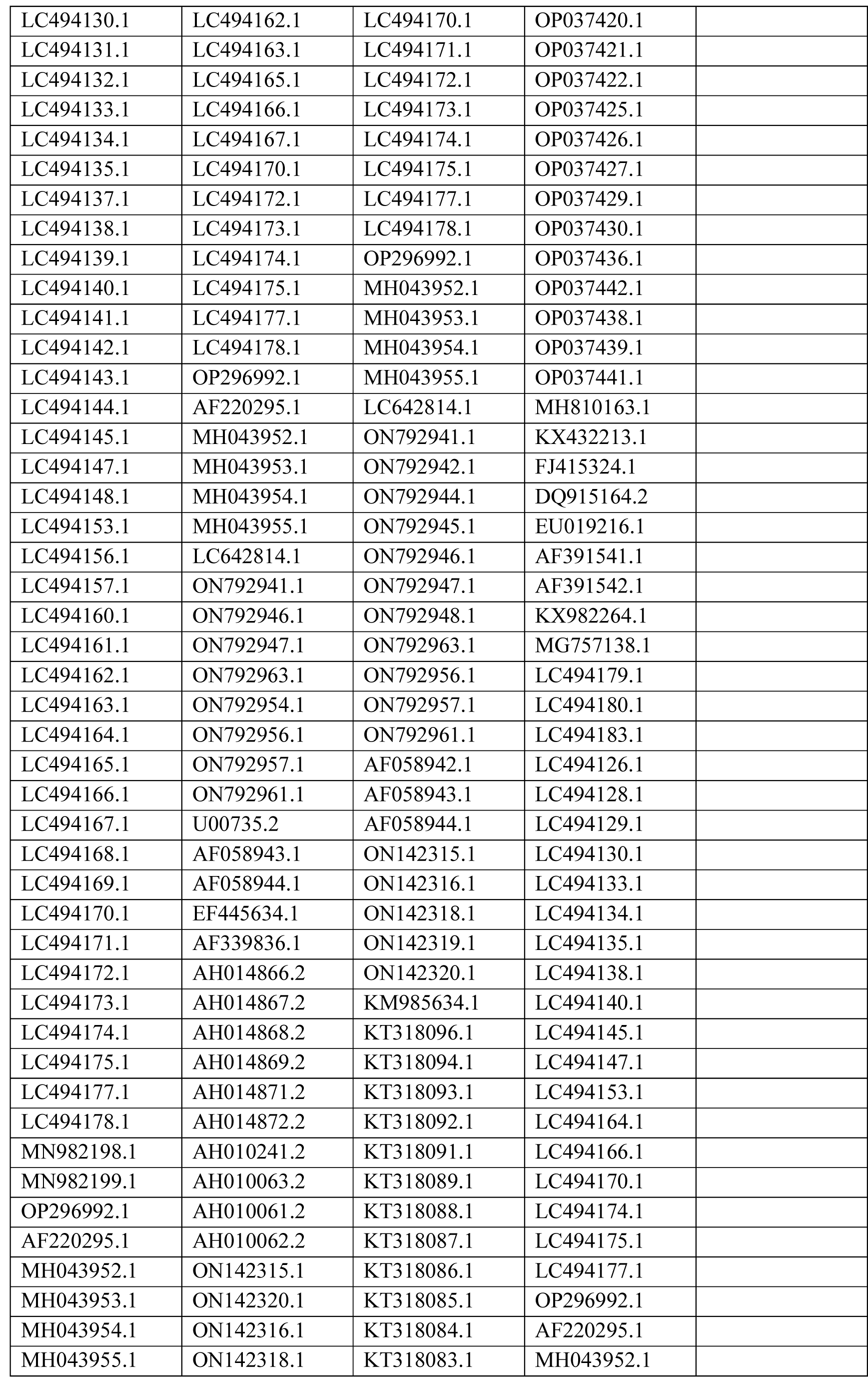
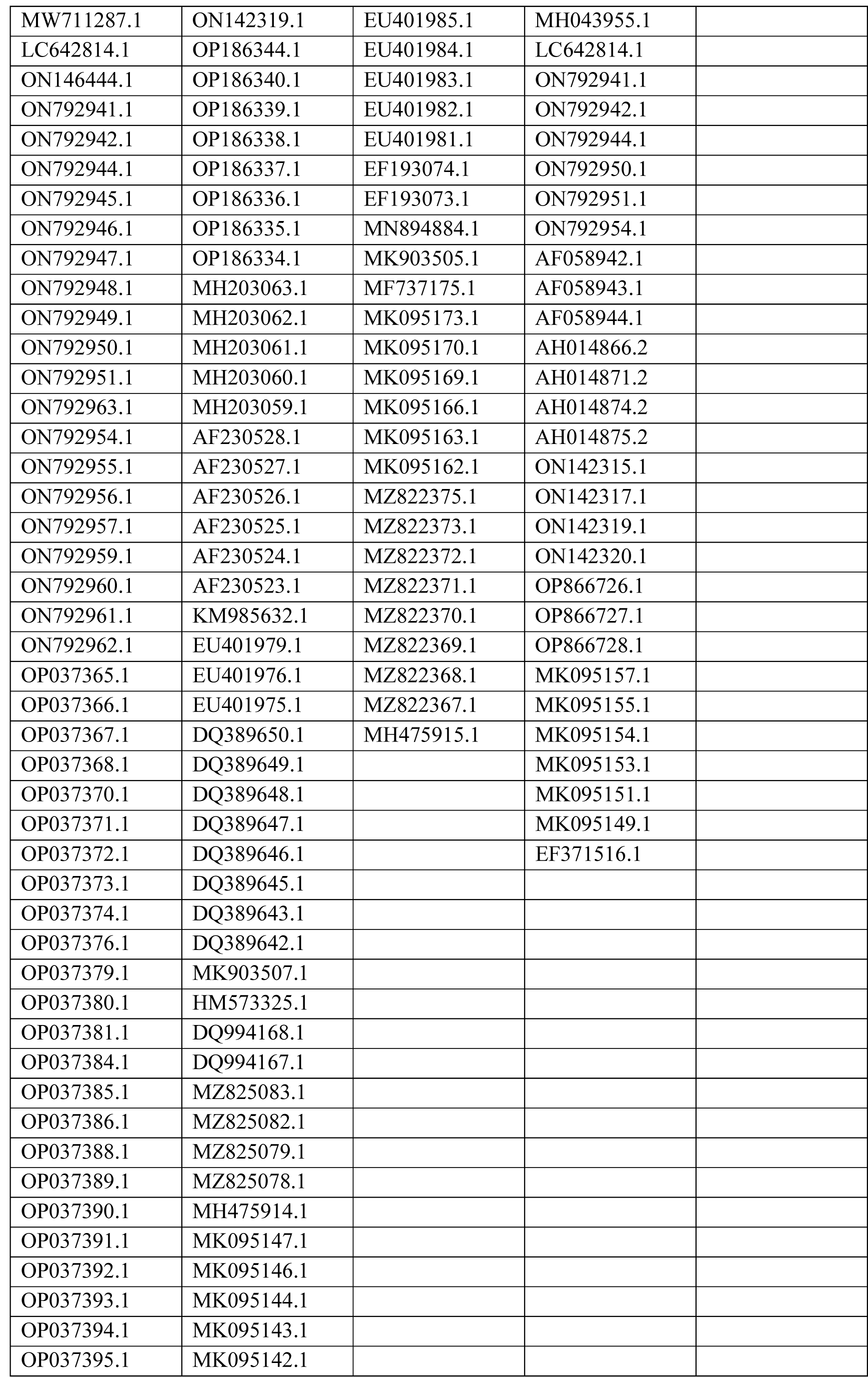
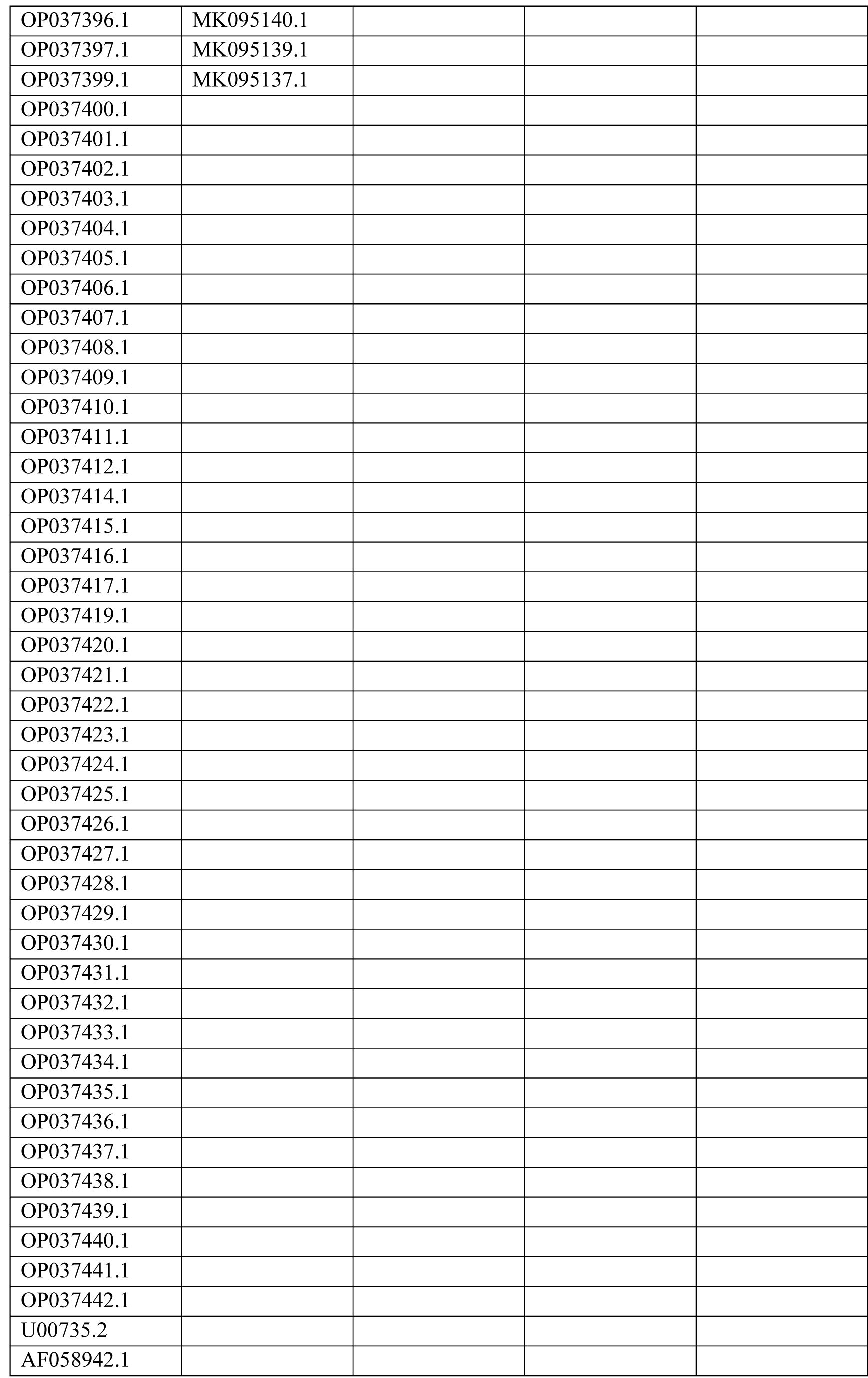
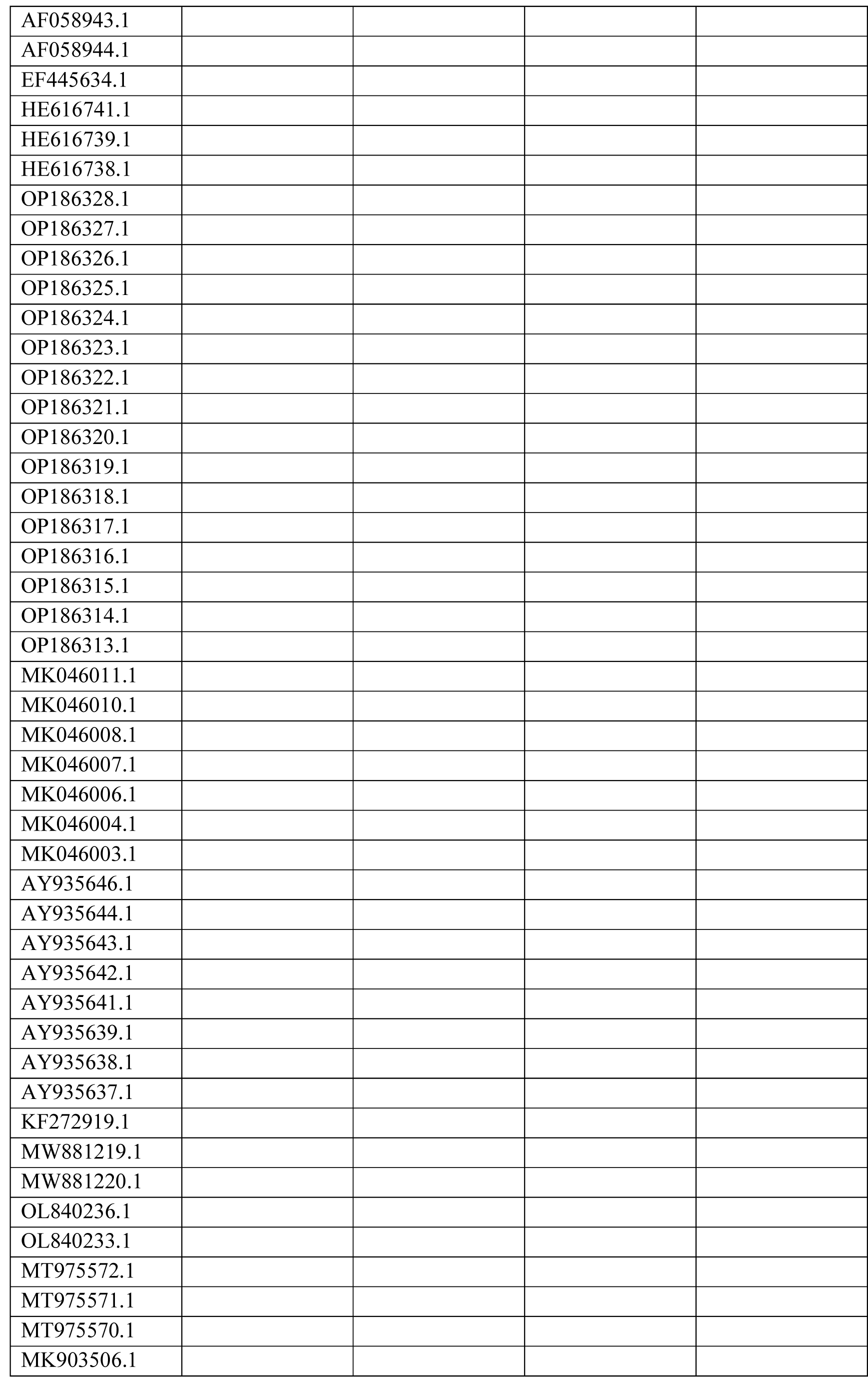
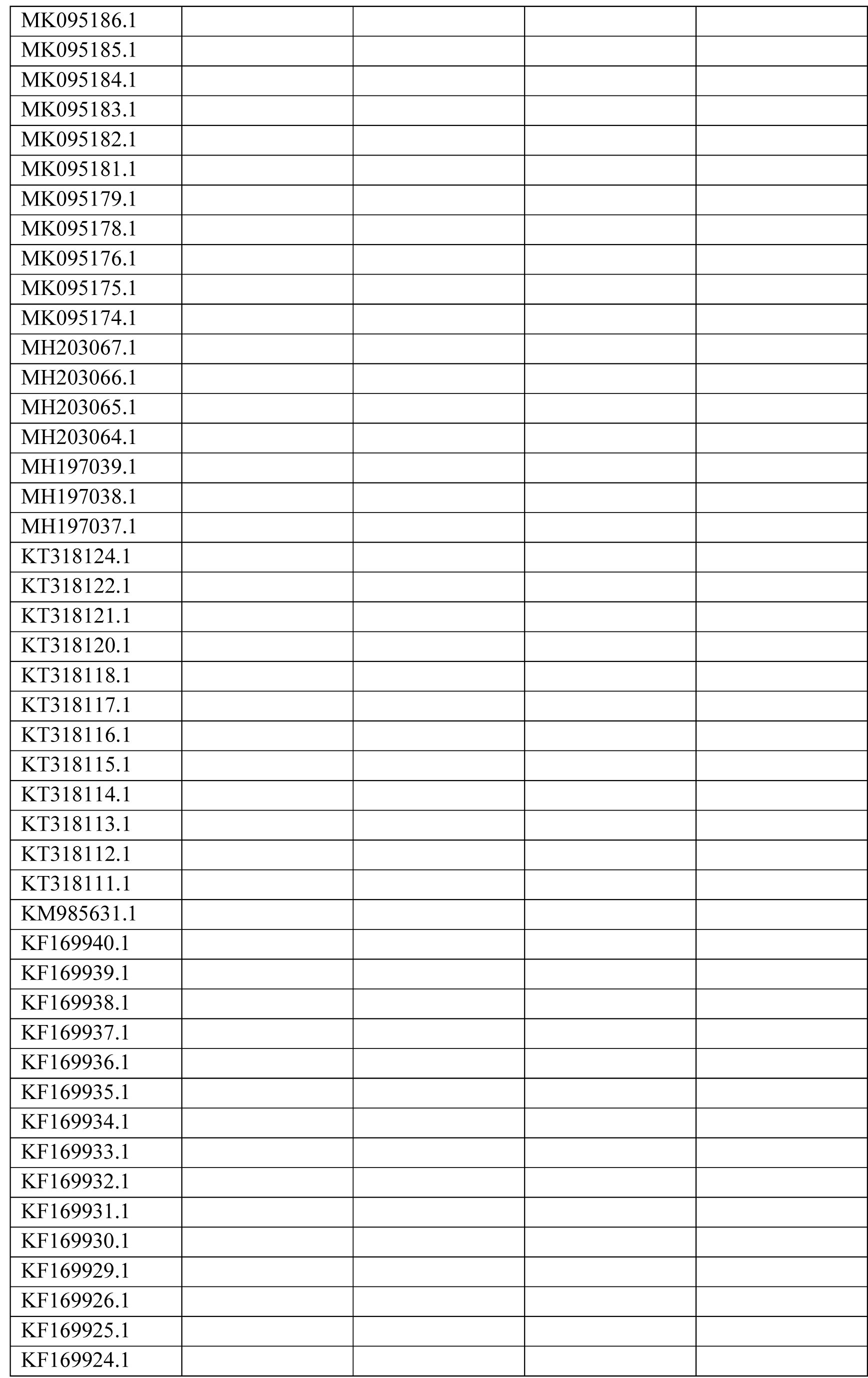
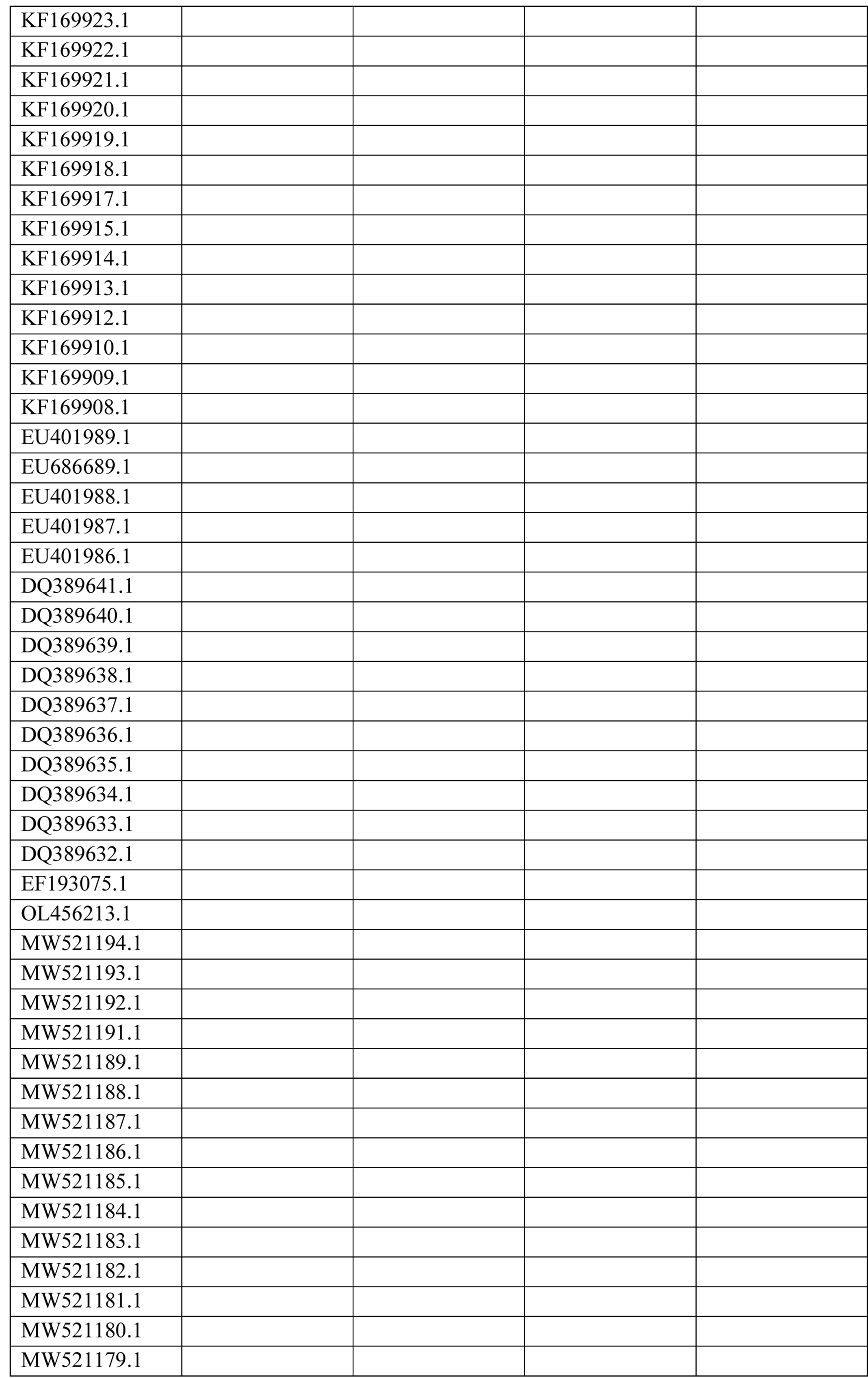
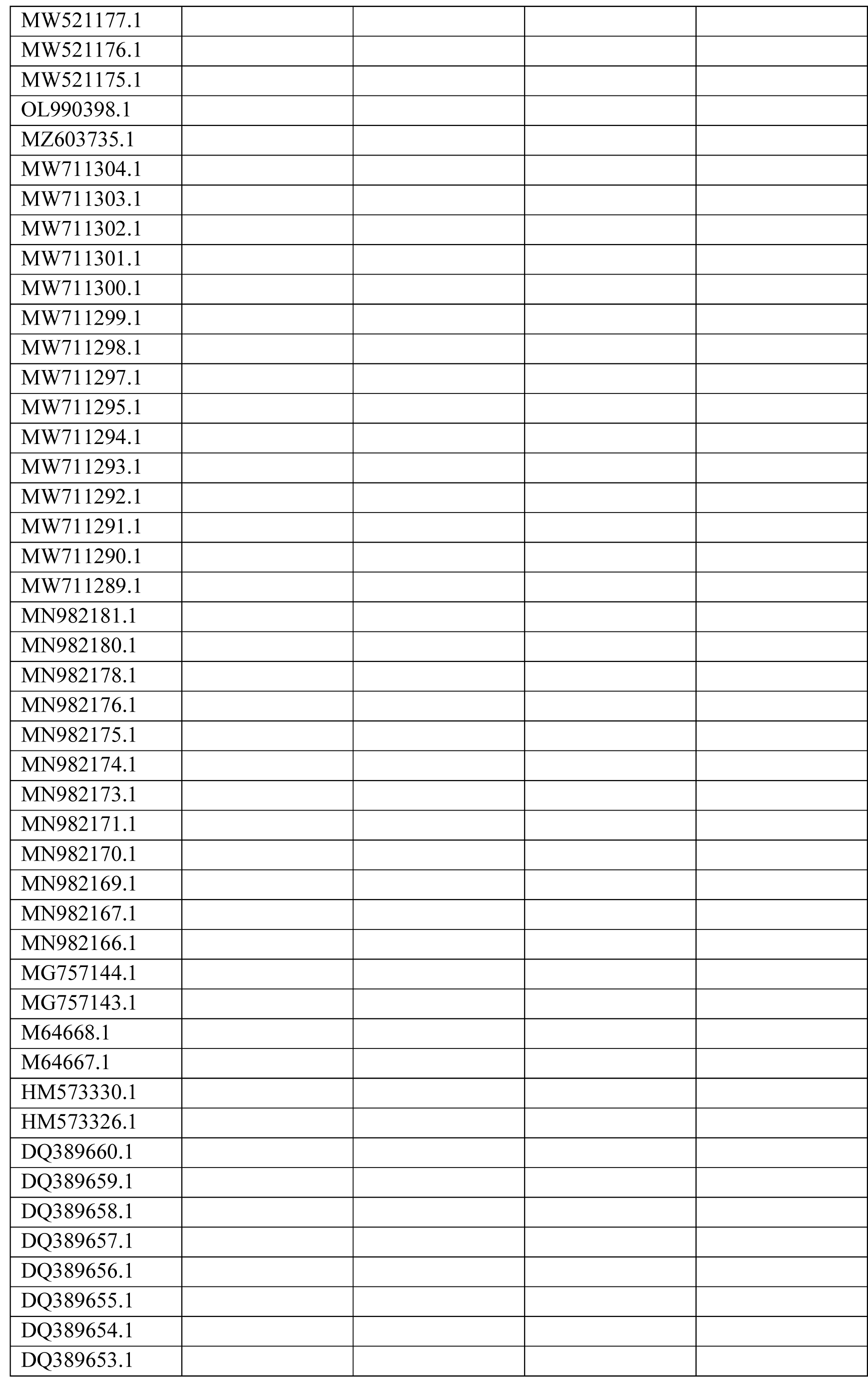
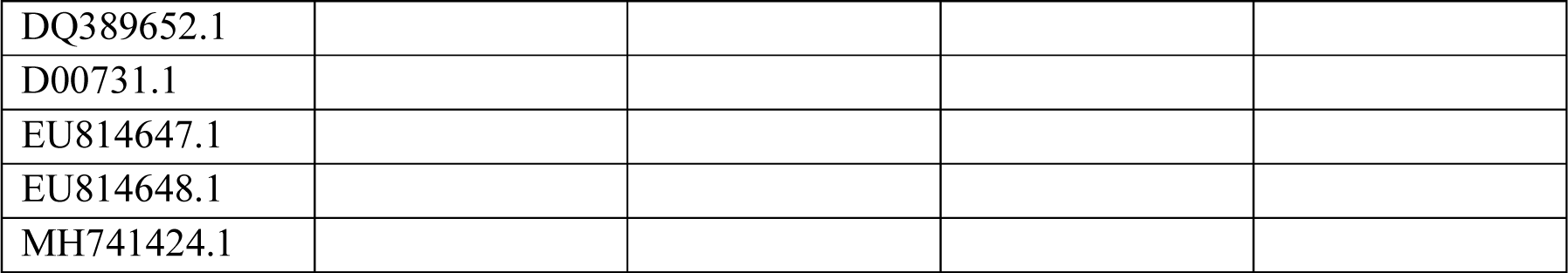
Spike, HE, nucleocapsid, membrane and small envelope GenBank Accession numbers for sequences used in selection and phylogenetic analyses.

## References

1. Vlasova AN, Saif LJ. Bovine Coronavirus and the Associated Diseases. Front Vet Sci 2021;8:293.

2. Gomez DE, Weese JS. Viral enteritis in calves. Can Vet J 2017;58:1267.

3. Fulton RW. Viruses in Bovine Respiratory Disease in North America: Knowledge Advances Using Genomic Testing. Vet Clin North Am Food Anim Pract 2020;36:321.

4. O’Neill R, Mooney J, Connaghan E, Furphy C, Graham DA. Patterns of detection of respiratory viruses in nasal swabs from calves in Ireland: a retrospective study. Vet Rec 2014;175:351–351.

5. Ellis J. What is the evidence that bovine coronavirus is a biologically significant respiratory pathogen in cattle? Can Vet J 2019;60:147.

6. Vlasak,’ R, Luytjes W, Leider,’ J, Spaan W, Palese1 P. The E3 protein of bovine coronavirus is a receptor-destroying enzyme with acetylesterase activity. J Virol 1988;62:4686–4690.

7. Lin X, O’Reilly KL, Burrell ML, Storz J. Infectivity-neutralizing and hemagglutinin- inhibiting antibody responses to respiratory coronavirus infections of cattle in pathogenesis of shipping fever pneumonia. Clin Diagn Lab Immunol 2001;8:357–362.

8. Milane G, Kourtesis AB, Dea S. Characterization of monoclonal antibodies to the hemagglutinin-esterase glycoprotein of a bovine coronavirus associated with winter dysentery and cross-reactivity to field isolates. J Clin Microbiol 1997;35:33–40.

9. Hussain KA, Storz J, Kousoulas KG. Comparison of bovine coronavirus (BCV) antigens: Monoclonal antibodies to the spike glycoprotein distinguish between vaccine and wild-type strains. Virology 1991;183:442.

10. Workman AM, Kuehn LA, McDaneld TG, Clawson ML, Chitko-McKown CG, et al. Evaluation of the effect of serum antibody abundance against bovine coronavirus on bovine coronavirus shedding and risk of respiratory tract disease in beef calves from birth through the first five weeks in a feedlot. Am J Vet Res 2017;78:1065–1076.

11. Workman AM, Kuehn LA, McDaneld TG, Clawson ML, Loy JD. Longitudinal study of humoral immunity to bovine coronavirus, virus shedding, and treatment for bovine respiratory disease in pre-weaned beef calves. BMC Vet Res 2019;15:1–15.

12. Dhufaigh KN, McCabe M, Cormican P, Cuevas-Gomez I, McGee M, et al. Genome Sequence of Bovine Coronavirus Variants from the Nasal Virome of Irish Beef Suckler and Pre- weaned Dairy Calves Clinically Diagnosed with Bovine Respiratory Disease. Microbiol Resour Announc;11. Epub ahead of print 17 November 2022. DOI: 10.1128/MRA.00821-22.

13. Gunn L, Collins PJ, O’Connell MJ, O’Shea H. Phylogenetic investigation of enteric bovine coronavirus in Ireland reveals partitioning between European and global strains. Ir Vet J;68. Epub ahead of print 30 December 2015. DOI: 10.1186/S13620-015-0060-3.

14. Decaro N, Elia G, Campolo M, Desario C, Mari V, et al. Detection of bovine coronavirus using a TaqMan-based real-time RT-PCR assay. J Virol Methods 2008;151:167.

15. Sievers F, Higgins DG. Clustal Omega for making accurate alignments of many protein sequences. Protein Sci 2018;27:135–145.

16. **Sievers** **F**, Wilm A, Dineen D, Gibson TJ, Karplus K, et al. Fast, scalable generation of high- quality protein multiple sequence alignments using Clustal Omega. Mol Syst Biol 2011;7:539.

17. Procter JB, Carstairs GM, Soares B, Mourão K, Ofoegbu TC, et al. Alignment of Biological Sequences with Jalview. Methods Mol Biol 2021;2231:203–224.

18. Waterhouse AM, Procter JB, Martin DMA, Clamp M, Barton GJ. Jalview Version 2—a multiple sequence alignment editor and analysis workbench. Bioinformatics 2009;25:1189–1191.

19. Troshin P V., Procter JB, Barton GJ. Java bioinformatics analysis web services for multiple sequence alignment—JABAWS:MSA. Bioinformatics 2011;27:2001–2002.

20. Kosakovsky Pond SL, Posada D, Gravenor MB, Woelk CH, Frost SDW. GARD: a genetic algorithm for recombination detection. Bioinformatics 2006;22:3096–3098.

21. Weaver S, Shank SD, Spielman SJ, Li M, Muse S V., et al. Datamonkey 2.0: A Modern Web Application for Characterizing Selective and Other Evolutionary Processes. Mol Biol Evol 2018;35:773–777.

22. Murrell B, Wertheim JO, Moola S, Weighill T, Scheffler K, et al. Detecting Individual Sites Subject to Episodic Diversifying Selection. PLOS Genet 2012;8:e1002764.

23. Murrell B, Moola S, Mabona A, Weighill T, Sheward D, et al. FUBAR: A Fast, Unconstrained Bayesian AppRoximation for Inferring Selection. Mol Biol Evol 2013;30:1196–1205.

24. Awadelkareem EA, Hamdoun S. Insilco Vaccine Design of spike and hemagglutinin esterase proteins of Bovine Coronavirus. Epub ahead of print 29 July 2022. DOI: 10.21203/RS.3.RS-1791707/V1.

25. Vijgen L, Keyaerts E, Moës E, Thoelen I, Wollants E, et al. Complete Genomic Sequence of Human Coronavirus OC43: Molecular Clock Analysis Suggests a Relatively Recent Zoonotic Coronavirus Transmission Event. J Virol 2005;79:1595.

26. Becerra-Artiles A, Nanaware PP, Muneeruddin K, Weaver GC, Shaffer SA, et al. Immunopeptidome profiling of human coronavirus OC43-infected cells identifies CD4 T cell epitopes specific to seasonal coronaviruses or cross-reactive with SARS-CoV-2. *bioRxiv*. Epub ahead of print 1 December 2022. DOI: 10.1101/2022.12.01.518643.

27. Wang C, Hesketh EL, Shamorkina TM, Li W, Franken PJ, et al. Antigenic structure of the human coronavirus OC43 spike reveals exposed and occluded neutralizing epitopes. Nat Commun;13. Epub ahead of print 1 December 2022. DOI: 10.1038/S41467-022-30658-0.

28. Waterhouse A, Bertoni M, Bienert S, Studer G, Tauriello G, et al. SWISS-MODEL: homology modelling of protein structures and complexes. Nucleic Acids Res 2018;46:W296.

29. Zeng Q, Langereis MA, Van Vliet ALW, Huizinga EG, De Groot RJ. Structure of coronavirus hemagglutinin-esterase offers insight into corona and influenza virus evolution. Proc Natl Acad Sci U S A 2008;105:90651.

30. 30. Schrödinger, L., & DeLano W. PyMOL. https://pymol.org/2/ (2020).

31. Hasoksuz M, Sreevatsan S, Cho KO, Hoet AE, Saif LJ. Molecular analysis of the S1 subunit of the spike glycoprotein of respiratory and enteric bovine coronavirus isolates. Virus Res 2002;84:101.

32. Martínez N, Brandão PE, de Souza SP, Barrera M, Santana N, et al. Molecular and phylogenetic analysis of bovine coronavirus based on the spike glycoprotein gene. Infect Genet Evol 2012;12:1870.

33. Saitou N, Nei M. The neighbor-joining method: a new method for reconstructing phylogenetic trees. Mol Biol Evol 1987;4:406–425.

34. Gouy M, Tannier E, Comte N, Parsons DP. Seaview Version 5: A Multiplatform Software for Multiple Sequence Alignment, Molecular Phylogenetic Analyses, and Tree Reconciliation. Methods Mol Biol 2021;2231:241–260.

35. Letunic I, Bork P. Interactive Tree Of Life (iTOL) v5: an online tool for phylogenetic tree display and annotation. Nucleic Acids Res 2021;49:W293–W296.

36. Franzo G, Drigo M, Legnardi M, Grassi L, Pasotto D, et al. Bovine Coronavirus: Variability, Evolution, and Dispersal Patterns of a No Longer Neglected Betacoronavirus. Viruses 2020, Vol 12, Page 1285 2020;12:1285.

37. Li X, Giorg EE, Marichannegowda MH, Foley B, Xiao C, et al. Emergence of SARS-CoV- 2 through recombination and strong purifying selection. Sci Adv;6. Epub ahead of print 1 July 2020. DOI: 10.1126/SCIADV.ABB9153/SUPPL_FILE/ABB9153_SM.PDF.

38. Dumonteil E, Herrera C. Polymorphism and Selection Pressure of SARS-CoV-2 Vaccine and Diagnostic Antigens: Implications for Immune Evasion and Serologic Diagnostic Performance. Pathog 2020, Vol 9, Page 584 2020;9:584.

39. Armijos-Jaramillo V, Yeager J, Muslin C, Perez-Castillo Y. SARS-CoV-2, an evolutionary perspective of interaction with human ACE2 reveals undiscovered amino acids necessary for complex stability. Evol Appl 2020;13:2168–2178.

40. Kistler KE, Bedford T. Evidence for adaptive evolution in the receptor-binding domain of seasonal coronaviruses OC43 and 229E. Elife 2021;10:1–35.

41. Zhang CY, Wei JF, He SN. Adaptive evolution of the spike gene of SARS coronavirus: Changes in positively selected sites in different epidemic groups. BMC Microbiol 2006;6:1–10.

42. Zhang F, Zang TM, Stevenson EM, Lei X, Copertino DC, et al. Inhibition of major histocompatibility complex-I antigen presentation by sarbecovirus ORF7a proteins. Proc Natl Acad Sci U S A;119. Epub ahead of print 11 October 2022. DOI: 10.1073/PNAS.2209042119/-/DCSUPPLEMENTAL.

43. Arshad N, Laurent-Rolle M, Ahmed WS, Hsu JC-C, Mitchell SM, et al. SARS-CoV-2 accessory proteins ORF7a and ORF3a use distinct mechanisms to downregulate MHC-I surface expression. bioRxiv 2022;2022.05.17.492198.

44. McGrath G, Tratalos JA, More SJ. A visual representation of cattle movement in Ireland during 2016. Ir Vet J 2018;71:1–5.

